# Injectable nanoclay gels for angiogenesis

**DOI:** 10.1101/566943

**Authors:** Daniel J. Page, Claire E. Clarkin, Raj Mani, Najeed A. Khan, Jonathan I. Dawson, Nicholas D. Evans

**Author notes:** Corresponding authors: Nicholas Evans and Jon Dawson.

## Abstract

The retention and sustained activity of therapeutic proteins at delivery sites are goals of regenerative medicine. Vascular endothelial growth factor (VEGF) has significant potential in promoting the growth and regeneration of blood vessels but is intrinsically labile. This is exacerbated by the inflammatory microenvironments at sites requiring regeneration. For VEGF to be efficacious it may require a carrier that stabilises it, protects it from degradation and retains it at a site of interest. In this study we tested the hypothesis that injectable nanoclay gels composed of Laponite XLG can stabilise VEGF and retain it in active form for therapeutic delivery. To achieve this, VEGF was incorporated in Laponite gels and its activity tested at a range of concentrations using *in vivo* cell culture tubulogenesis assays and *in vivo* angiogenesis assays. We found that VEGF-Laponite gels enhanced tubulogenesis in a dose-dependent manner *in vivo*. When administered subcutaneously *in vivo* Laponite was retained at an injection site for up to a period of three weeks and promoted a 4-fold increase in blood vessel formation compared with alginate or vehicle controls as confirmed by CD31 staining. Notably, in contrast to alginate, Laponite gels did not release VEGF, indicating a strong interaction between the growth factor and the nanoclay, and suggesting that Laponite enhancement of VEGF efficacy is due to its retention at an implantation site over a prolonged period. Our approach provides a robust method for delivery of bioactive recombinant VEGF without the necessity for complex hydrogel or protein engineering.

## Introduction

Preservation and retention of the activity of therapeutic proteins at the site of delivery are major goals in regenerative medicine. Growth factors, such as vascular endothelial growth factors (VEGF), platelet-derived growth factors (PDGF) and fibroblast growth factors (FGF) have significant potential in promoting the growth and regeneration of cells and tissues [1]. However, proteins are intrinsically labile and are often not long-lived enough to ensure significant biological effect [2]. This may be compounded in sites requiring regeneration, which are often characterised by inflammation and high local concentrations of proteases. Furthermore, local injection of soluble growth factors may only result in transient activity before systemic clearance via the lymphatic and circulatory systems [3]. For these reasons, it is likely that for growth factors to be efficacious they require a carrier that stabilises and protects a growth factor at a site of interest.

VEGFs comprise a family of growth factors that are responsible for increasing vascular permeability, endothelial cell signalling and proliferation to stimulate angiogenesis [4,5]. Recombinant VEGF proteins, most notably the predominant form of VEGF-A, VEGF_165_, have been investigated for more than 20 years as a therapy for promoting angiogenesis in degenerative disease or injury [6], to treat for example, stroke, bone fracture and skin wounds. However, their use is limited by the high doses required [7], subsequent off-target effects such as hypotension [8] and suboptimal angiogenesis causing, for example, severe vascular leakage and subsequent oedema [9].

To overcome these challenges, VEGFs can be incorporated within solid scaffolds, either naturally-derived or synthetic, to control release. Early studies used alginate as a depot to deliver active VEGF to cultured cells [10]. This seaweed-derived polysaccharide material provides a relatively inert carrier scaffold that can be gelled in situ using a divalent cation such as Ca^2+^ but which does not significantly restrict protein diffusion. To better control release, scaffold physical properties may be modified – for example by controlling surface area to volume ratio of polymer microparticles [11]. Alternatively, growth factors have been covalently crosslinked to the scaffold ‘backbone’. Zisch *et al.* demonstrated enhanced angiogenesis when VEGF was covalently crosslinked with donor-derived fibrin [12]. Some biomaterial strategies have taken advantage of the affinity of heparin for VEGF by incorporating heparin within backbone of the scaffold, so promoting VEGF retention [13–15], while VEGF has also been covalently linked directly to the backbone of polymeric hydrogels (based on PEG [16]). An advantage of these biomaterials is that other peptide motifs can be engineered into the polymeric backbone. For example, MMP-sensitive peptides or cell binding motifs, such as peptides containing the RGD amino acid motif. Together this allows concurrent cell attachment ingrowth, degradation and growth factor release. These biomaterials have found application in a wide range of preclinical models, such as in cardiac repair [17] and stroke [18], but their translation to the clinic is limited by their complexity. There remains a pressing need for simple biomaterials that better localise and preserve growth factor activity.

We are investigating the notion that inorganic, clay nanomaterials may provide an answer to these shortcomings. Clays may be loosely defined as materials that comprise finely-grained inorganic particles that, when mixed with water, form colloidal pastes or gels [19]. Clays are highly sorptive for biomolecules, a property which has been exploited in their use over many years as orally-taken gastrointestinal protectors [20], skin exfoliation agents [21,22] and as blood clotting agents [23]. More recently we [24,25] and others [26,27] have tested the idea that synthetic nanoclays, such as Laponite XLG (Na^+^_0.7_[(Mg_5.5_Li_0.3_Si_8_O_20_(OH)_4_]^−^_0.7_,) may be suitable as delivery agents of proteins for regenerative medicine.

Like many clays, Laponite has a long history of use in the food and cosmetic industries, suggesting that it is non-toxic and well tolerated, even at high doses. Aqueous dispersions of Laponite-mineral nanoplatelets (25 nm in diameter and ∼1 nm in thickness) display self-assembly properties due to their charge anisotropy and form thixotropic (shear-thinning), clear, colloidal gels. These properties allow injection through a hypodermic needle and the spontaneous formation of stiff, irreversible gels upon contact with blood proteins and ions through a diffusion gelation mechanism, without the need for further chemical modification [28]. These properties make clays extremely attractive as, for example, tissue fillers for regenerative medicine, or in dressings or salves at the skin surface.

In this study we tested the hypothesis that the activity of VEGF_165_ could be preserved and sustain an angiogenic response following incorporation within the bulk of injected Laponite gels.

## Methods

### Biomaterial preparation

3% (w/v) suspensions of Laponite XLG (BYK Additives, Widnes, UK) were prepared by slowly adding Laponite powder to distilled water under rapid agitation. Suspensions were autoclaved and volume adjusted with sterile ddH_2_O. 1.1% (w/v) alginate solutions were prepared from anhydrous ultra-pure alginate (NovaMatrix, Sandvika, Norway), with UV sterilisation for 30-60 minutes prior to preparation. To crosslink the alginate gel suspensions, CaCl_2_ was added at a final concentration of 100 mM.

### Human umbilical vein endothelial cell (HUVEC) isolation and culture

HUVECs were isolated from umbilical cords collected from the Princess Anne Hospital, Southampton, UK, with the approval of Southampton and South West Hampshire Local Research Ethics Committee (Ref:05/Q1702/102). HUVECs were isolated and cultured as described Dawson *et al* [24] with some minor modifications. Briefly, the umbilical cord was cut at both ends; to one end, a cannula was inserted into an exposed vein and secured with ties. Sterile PBS was first flushed through the cord until the waste PBS collected at the other end was clear. The umbilical cord clamped at one end and sufficient 5 mg/ml collagenase B (Sigma-Aldrich, Poole, UK) added to fill the vein before incubation at room temperature for 1 hour. The collagenase solution was then removed, centrifuged at 200*g* for 5 minutes and the supernatant discarded. Cells were re-suspended in endothelial cell growth medium (ECGM), which consisted of: Medium 199 (Lonza), 10% fetal bovine serum (FBS; Life Technologies, Paisley, UK; product: 10270106, batch: 41Q4297P), penicillin-100U/ml-streptomycin-100U/ml (Sigma-Aldrich/Merck) and endothelial cell growth supplement (Promocell GmbH, Heidelberg, Germany). Re-suspended cells were cultured at 37°C/5% CO_2_ in humidified conditions. Following incubation, cells were sub-cultured using ECGM and passages of 1-4 were used in all experiments.

### Release of VEGF by Laponite/alginate hydrogels

To assess the release of VEGF from biomaterials, VEGF was premixed with by Laponite or alginate at a concentration of 40 µg/ml. 10 µl aliquots of biomaterial-VEGF were transferred into low-protein binding tubes (Eppendorf® LoBind) containing 90 µl assay diluent. Biomaterials containing no VEGF and media with aqueous VEGF added served as negative and positive controls respectively. Media was incubated at 37 C /5% CO_2_ in humidified conditions and then recovered at various time points up to 3 weeks and stored at −20 □ until required for protein analysis.

To analyse the protein content from the supernatant recovered from biomaterial tubes, an enzyme-linked immunosorbent assay (ELISA) (R&D Systems, Abingdon, UK) was performed according to the manufacturer’s instructions.

### HUVEC 2D tubule formation on Laponite gels

To promote cell attachment and growth on Laponite gels, human fibronectin (Merck-Millipore, Watford, UK) was premixed with Laponite gels to a final concentration of 50 µg/mL. Cell culture substrates were made by adding 150 µl of Laponite to the bases of wells of 24-well plates, before briefly centrifuging to ensure an even Laponite layer and incubation for 1 hour at 37 C for gelation.

To test the activity of VEGF_165_ (PeproTech, London, UK; hereafter denoted VEGF) in promoting tubule formation when associated with the Laponite biomaterials, VEGF was either mixed or surface adsorbed to pre-formed Laponite gels. For the former, reconstituted human recombinant VEGF was added to final concentrations of 1 to 5 µg/ml and mixed by vortexing prior to forming substrates. For the latter, VEGF in ECGM was incubated at a concentration of 12.6 ng/cm^2^ at room temperature for one hour.

Following formation of substrates, HUVECs were re-suspended in ECGM containing 10% FBS and 40 ng/ml basic fibroblastic growth factor (bFGF; Thermo Fisher Scientific-Invitrogen, UK) and seeded at 5 × 10^4^ cells/well (n=6). To test the activity of soluble growth factor, VEGF was added to the medium at concentrations of 0 – 40 ng/mL.

### Image analysis of HUVEC 2D tubule network

A freely available macro for ImageJ called ‘Angiogenesis Analyzer’ by Gilles Carpentier (Gilles Carpentier. Contribution: Angiogenesis Analyzer, ImageJ News, 5 October 2012, http://image.bio.methods.free.fr/ImageJ/?Angiogenesis-Analyzer-for-ImageJ#nb1) for NIH Image J 1.47v Program was used to automatically quantify HUVEC 2D tubule network. Please refer to Supplementary Information 1 [S1] for more information regarding software configuration settings.

### Animals

18 male MF1 mice (Biomedical Research Facility, University of Southampton) aged between 8-10 weeks old were used to investigate localisation of VEGF using Laponite hydrogels. All animals were bred and maintained at the Biomedical Research Facility, Southampton in a temperature-controlled environment (20 C-22 C) with a 12 h light/dark cycle (lights on at 0600 hrs). Food and water were provided ad libitum both pre- and post-injection. All work was done under Ethical Approval obtained under project licence PPL 30/2971.

### Subcutaneous VEGF-biomaterial treatments

Mice were anesthetised using inhaled anaesthetic (isoflurane; product from Centaur, Castle Cary, UK) and the whole dorsal region shaved. On the left side of the dorsum, pre-prepared Laponite gels were administered subcutaneously by injection (50 µL) at three separate locations (rostral to caudal) with three different VEGF doses (0.1, 1.0 and 4.0 µg total VEGF) (n = 6 mice). On the contralateral side 3 vehicle Laponite treatments were administered (negative control). A control biomaterial hydrogel, alginate was administered in the same way (n = 3 mice). Mice were killed by CO_2_ asphyxiation for 3-5 minutes followed by cervical dislocation. Biomaterial and surrounding cutaneous tissue was surgically removed after 21 days and fixed in 4% (w/v) paraformaldehyde (PFA) for 18 hours. Tissue samples were then transferred to 70% ethanol and stored at 4°C until required for processing.

### Unbiased macroscopic angiogenic scoring

Upon tissue harvest, photographs of each biomaterial treatment and dose were captured using a Nikon D3200 digital single-lens reflex (SLR) camera. A scaled ruler was present for all images captured. All images were then labelled (biomaterial treatment, VEGF concentration, time point). Using the Measure module within ImageJ, the assigned label was ‘measured’ using the Batch command to generate a list of all the image names with a corresponding number. These data were imported into Microsoft Excel (2016 version). Within Excel, a column was inserted adjacent to the label data set. In the first cell of this column the =RAND() command was entered, and then copied into every cell; this command created a list of random numbers which then allowed the label data set to be sorted to this random list. The images were then arranged on blank page (no label included). The sheets that contained these randomised images were used as the basis of a randomised blinded questionnaire used to measure the degree in which angiogenesis had occurred. Please refer to Supplementary Information 2 [S2] for a copy of the questionnaire that was designed for this study.

### Histological analysis

Fixed samples were washed in PBS prior to incubation with 30% (w/v) sucrose (Sigma-Aldrich/Merck) in PBS overnight at 4 C. The samples were then briefly washed in optimal cutting temperature compound (OCT) (CellPath, Newtown, UK) to remove excess sucrose and then immersed with OCT in a cryo-mould. Cryo-moulds were placed into a solution of pre-cooled (−80°C) isopropanol (dry ice was added to isopropanol) to allow controlled sample freezing. Moulds were stored at −80°C until required for sectioning. Sequential sections were cut at 10 µm thickness using a cryotome (maintained at −25 C to −30 C) and mounted on charged glass slides and placed on a warming rack (37 C) for 30 minutes and then stored at − 80 C. When required for histological and immunohistological staining, cryo-sections were thawed at room temperature for 10 minutes and washed in PBS for 10 minutes.

### Histochemistry

#### Anti-CD31 immunohistochemistry

A rat detection kit for anti-mouse CD31 (MenaPath, Menarini Diagnostics, Winnersh, UK; product: MP-517-RTK6) was used in combination with an anti-CD31 antibody (rat monoclonal, MenaPath, Menarini Diagnostics, product: MP-303-CM01) following manufacturer’s instructions. Slides were dehydrated and mounted with cover slips using DPX mounting medium (Thermo Fisher; product: 10050080).

#### H&E

Following OCT removal, slides were stained with hematoxylin and eosin before mounting with coverslips

#### Auramine O

Slides were flooded with pre-prepared Auramine O (Auramine O 0.3 g, phenol 3.0 g, distilled water 100 ml) (Sigma-Aldrich/Merck) for 15 minutes at room temperature in the dark. Excess Auramine O solution was discarded and slides washed in distilled water (2 × 2-minute washes). Slides were immediately mounted with coverslips using Fluoromount™ (Sigma-Aldrich/Merck; product: F4680-25ML) and sealed with nail varnish. Slides were stored in the dark and imaged within 3 days.

### Microscopy

Histological sections were imaged using the Olympus BX 51 dotSlide virtual slide microscope system (Olympus Life Science, Southend-on-Sea, UK). Captured images were extracted and analysed using Fuji ImageJ v1.50b, BIOP PT VSI plugin and Olympus Virtual Slide Desktop 2.4 software.

### Analysis of CD31 staining by Chalkley count

The Chalkley point-overlap morphometric technique (more simply referred to as “Chalkley method”) is a relative area estimate method to measure the abundance of microvessels in a immunohistochemical sample [29]. Adapting a similar approach documented in a previous publication [30], a digital Chalkley grid overlay was applied to a x20 magnification image at 3-5 different “hot-spot” regions. Counts of these 3-5 regions that landed on the most positively stained structures through rotating the digital overlay were recorded and the mean values used for analysis.

### Analysis of cellularity

H&E-stained image samples were imported into a free open-source software program called Orbit Image Analysis (revision 2.67; http://www.orbit.bio/download/). This software was used to isolate/segment haematoxylin-stained (nucleated) cells and measure and compare the % area of pixels. Please refer to Supplementary Information 3 [S3] for software configuration settings.

### Statistical analysis

A one-way analysis of variance (ANOVA) (parametric) was used for statistical analysis of blood glucose level data and mouse weight data Tukey’s multiple comparison test was used to determine individual *p* values. A two-way ANOVA (parametric) was used for statistical analysis of wound closure rate data and Tukey’s multiple comparison test was used to determine individual p values. A student’s t-test (unpaired) was used for statistical analysis of CD31 staining data.

## Results

### VEGF is sequestered tightly by Laponite gels and is not released

To test the release profile of VEGF premixed within an injectable Laponite gel formulation, we performed an *in vivo* release assay using an ELISA for VEGF. Over a period of 21 days, no VEGF was released from Laponite-VEGF gels (incorporated at a concentration of 40 μg/mL). In direct contrast, VEGF was released rapidly from alginate-VEGF gels, with 43.1 ± 4.3% released at 12 hours and 77.4 ± 6.6% & at 24 hours (Figure 1A).

**Figure 1.**
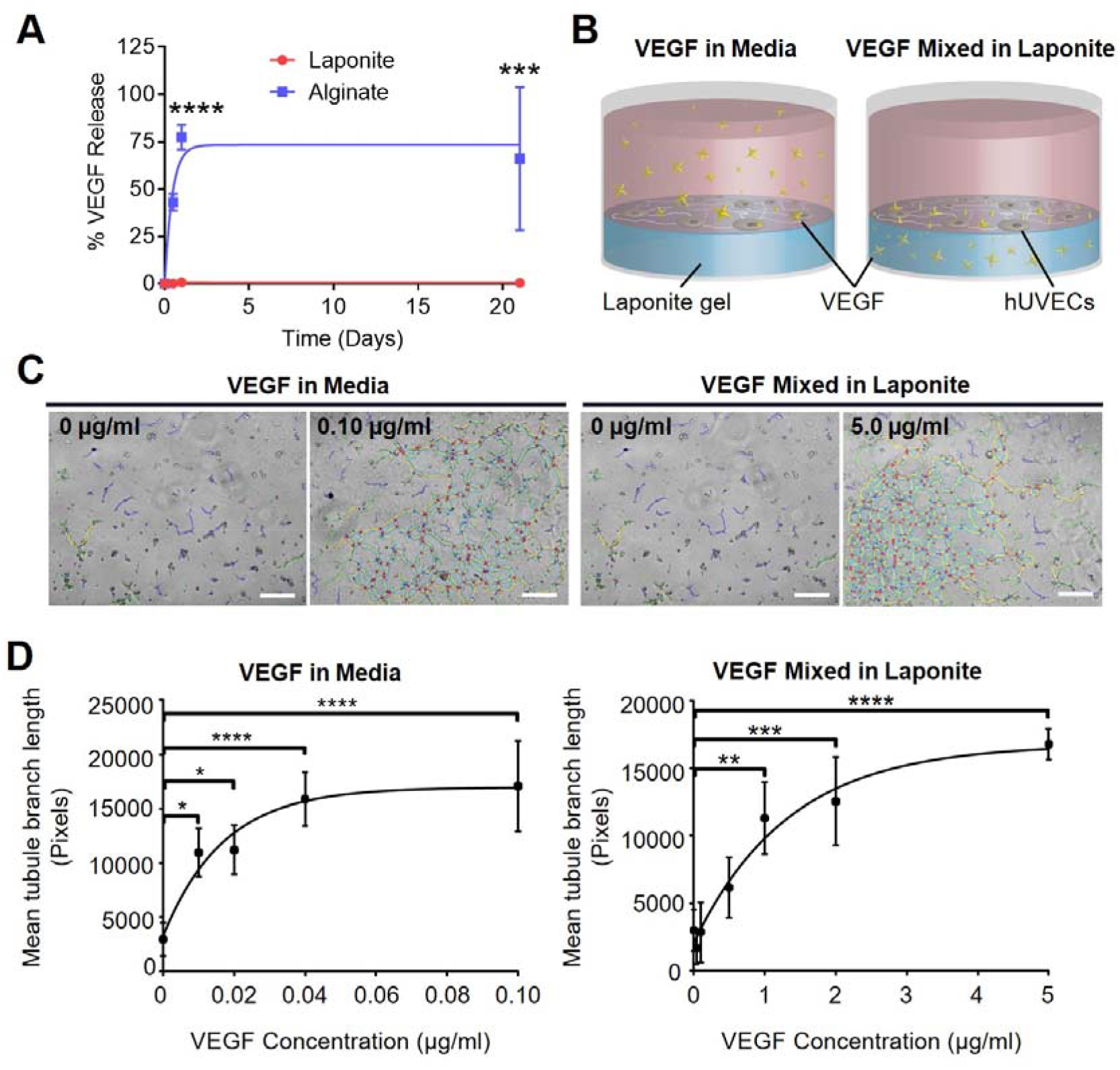
Localisation of VEGF by Laponite hydrogels stimulates in vivo angiogenesis. (A) VEGF was not released from Laponite gels over a period of 21 days (red plot). This contrasted with alginate gels, where ∼75% of incorporated VEGF was rapidly released over 24 hours. *** p <0.001; **** <0.0001. n = 6; error bars indicate SEM. (B) Schematic illustrating 2D tubule formation of human vein endothelial cells (HUVECs) on Laponite hydrogels study design. Left-hand image shows schematic VEGF in media, and right-hand image shows incorporation of VEGF_165_ incorporated in Laponite gel (VEGF illustrated with yellow stars). (C) VEGF stimulates tubule formation in HUVECs cultured with VEGF present in medium (left) or mixed in Laponite (right), as shown by phase contrast microscopy with network formation analysis segmentation (scale bars = 500 µm). (D) Network formation in response to VEGF was dose-dependent, but with higher concentrations required when mixed in Laponite compared with added to the medium. Error bars = SD, n = 3, one-way ANOVA analysis performed with Tukey’s post-hoc test applied for multiple comparisons; * p < 0.05 ** p < 0.01 ** p < 0.001 ****p < 0.0001.

### Localisation of VEGF by Laponite stimulates angiogenesis in vivo

VEGF has previously been shown to retain bioactivity following surface-adsorption to Laponite nanoclay from aqueous media [24]. Its utility would be substantially increased, however, if it could be delivered stably within the bulk of a gel or paste, for example as an injectable or topically applied material. To explore the potential of this approach, we first tested whether the activity of VEGF was preserved following incorporation into the bulk of Laponite gels in comparison to aqueous VEGF. HUVECs, which form quantifiable tubule networks in response to VEGF, were cultured on the surface of 3% (w/v) Laponite gels pre-mixed by vortexing with VEGF at concentrations between 0 and 5.00 µg/ml. For comparison cells were cultured on the surface of 3% Laponite gels (premixed with vehicle alone; no VEGF) in the presence of aqueous VEGF at concentrations 0 – 40 ng/ml (a schematic of the experimental design is shown in Figure 1B).

Aqueous VEGF promoted HUVEC tubule formation on Laponite substrates in a dose-dependent manner, with a maximal effect at a concentration of 0.04 μg/mL and a half-maximal effect at ∼0.01 μg/mL (Figure 1C). VEGF adsorbed at a concentration of 12 ng/cm^2^ also promoted maximal tubule formation (Supplementary Figure 1) as demonstrated in a previous publication [31]. When VEGF was mixed in the bulk of the Laponite, it retained activity. However, a concentration of 5 µg/mL VEGF was required to exhibit the equivalent stimulatory effect as seen with aqueous VEGF and Laponite-adsorbed groups, with a half maximal response at ∼ 1 µg/mL. (p = <0.01) (Figure 1C). Together these data indicate that VEGF activity is preserved following following mixing in the bulk of the Laponite, similarly to surface-adsorption, but that the latter reduces either bioavailability or protein activity.

### Laponite-incorporated VEGF stimulates and supports blood vessel infiltration

We next tested whether VEGF mixed in 3% (w/v) Laponite (hereafter referred to as VEGF-Laponite) could induce angiogenesis *in vivo*. VEGF-Laponite was injected subcutaneously on the dorsa of male aged-matched mice (Figure 2A). Based on our *in vivo* observations, we chose a range of VEGF concentrations between 1 and 40 µg/ml (total doses of between 0.05 µg and 2 µg). Alginate, a well-understood biomaterial used in drug delivery [32] was used as a control hydrogel at a concentration of 1.1 % (w/v) – chosen for its similar rheological profile – containing the same range of VEGF concentrations.

**Figure 2.**
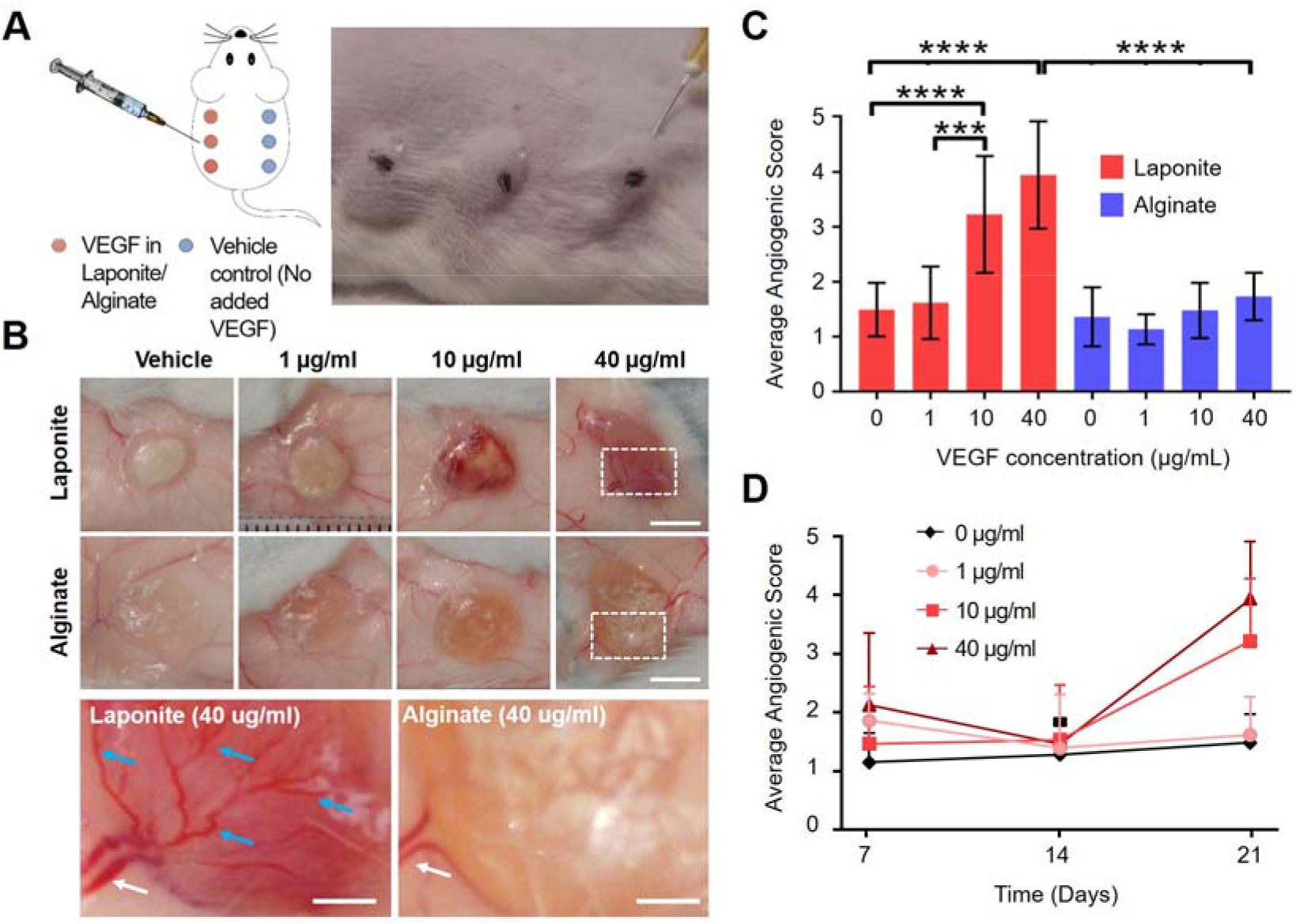
VEGF encapsulated by Laponite hydrogels stimulates in vivo angiogenesis. (A) A schematic showing subcutaneous injection of Laponite or alginate on the dorsum of healthy mice at a range of concentrations. A vehicle control was administered on the contralateral side. (B) Increased angiogenesis was visible in macroscopic images at biomaterial sites that contained high VEGF (> 1 µg/ml) concentrations. Blood vessels were observed infiltrating the Laponite and branching over the surface (blue arrows shows vessel branching, white arrows shows vessels located at the biomaterial periphery) in VEGF-Laponite hydrogels. Scale = 5 mm. (C) Blinded scoring of harvested biomaterials confirmed that there was a significant increase in angiogenesis in Laponite-VEGF treated groups at VEGF concentrations 10 and 40 µg/ml at day 21 versus alginate-VEGF. (D) Differences in blood vessel infiltration only became apparent at day 21; no significant change in anngiogenic score was present at day 14. n = 3; Error bars = SD. Two-way ANOVA analysis performed with Tukey’s post-hoc test applied for multiple comparisons. * p < 0.05 ** p < 0.01 ** p < 0.001 ****p < 0.0001.

Laponite and alginate biomaterials could be observed macroscopically at injection sites for at least 21 days (Figure 2B). At tissue harvest, blood vessel infiltration of VEGF-Laponite gels at concentrations of 10 or 40 µg/mL was evident (Figure 2B). At 40 µg/mL blood vessels from the surrounding tissue were observed growing into and along the surface of the implanted VEGF-Laponite. In some cases, extensive vessel branching was present (Figure 2B lower panel; arrows). In contrast, alginate-VEGF hydrogels exhibited less overt ingrowth. Even when 40 µg/mL VEGF was incorporated, pre-existing blood vessels present at the periphery of the alginate implant (Figure 2B lower panel) showed little evidence of branching into the biomaterial.

To quantify the degree of blood vessel infiltration, blinded observers scored images of samples on a scale of 0 – 6 (see Supplementary Information 2 questionnaire for details of scoring criteria). Blood vessel infiltration scored significantly greater in Laponite gels delivered at concentrations of ≥ 10 μg/mL compared to ≤ 1 μg/mL (n = 6, *p* < 0.001) (Figure 2C). In contrast, no significant increase in blood vessel infiltration was measured in alginate-VEGF gels at any concentration. The increase in vessel infiltration was time-dependent; there were no significant changes in blood vessel infiltration for any gel at d14, with increases only becoming evident at d21 (Figure 2D). These data indicate that, in contrast to alginate, Laponite preserves the activity of VEGF *in vivo*, either by preventing release of the growth factor or by preserving its activity.

### Laponite-incorporated VEGF stimulates angiogenesis and cell invasion in vivo

Next, we investigated cell infiltration and angiogenesis in implanted VEGF-containing Laponite and alginate gels. Harvested biomaterial tissue samples were sectioned and histological staining performed. Staining of tissue sections with anti-CD31 (an endothelial cell marker) showed greater density of blood vessels at higher (10 & 40 µg/ml) VEGF concentrations when localised by Laponite gels; in contrast, negligible CD31 was apparent within the alginate biomaterial treatment (Figure 3A). Quantification of CD31+ blood vessels was performed using a Chalkley grid analysis (Figure 3B) which confirmed an increase in number of vessels with increasing VEGF concentration in Laponite samples (however only 40 µg/ml VEGF was shown to be significantly different when compare to other concentrations and Laponite vehicle (*p* = <0.01)). In summary these data indicate that injectable Laponite gel facilitates the ingress of cells and blood vessels, and that this is promoted by the inclusion of VEGF in the bulk of the gel.

**Figure 3.**
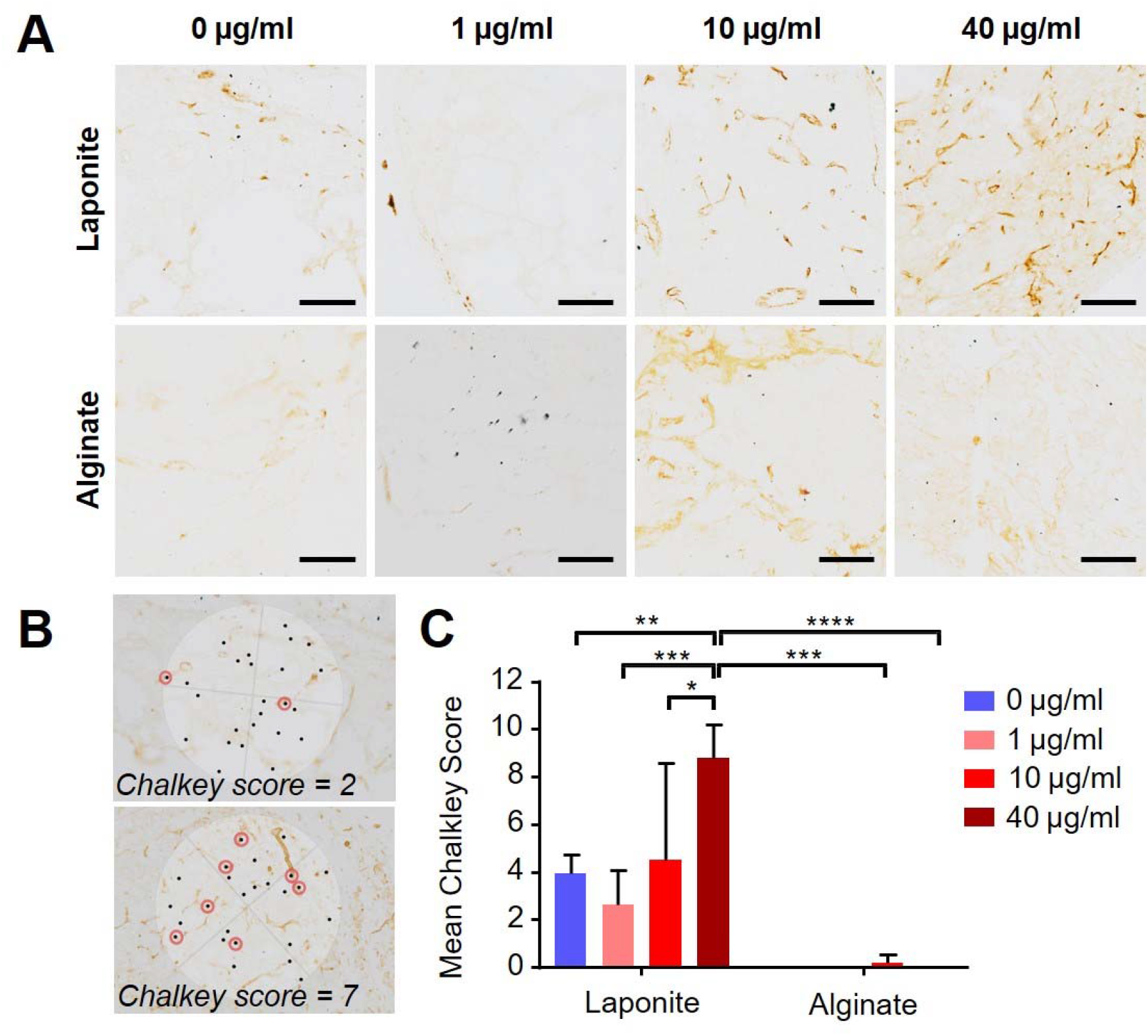
Localisation of VEGF encapsulated by Laponite hydrogels stimulates in vivo angiogenesis; immunohistochemical analysis. (A) The density of blood vessels was markedly greater in VEGF-Laponite compared to Laponite alone, as indicated by anti-CD31 staining. This was not obvious in VEGF-alginate controls. Scale = 100 µm. (B) Blood vessel density was quantified using a Chalkley grid (see methods); the upper panel is an example of a low Chalkley count (2 vessels identified) and the lower panel an example of a high count (7 vessels identified) (shown by the red circles) (C) Chalkley count analysis showed that 40 µg/ml VEGF mixed within Laponite caused significant growth of novel blood vessels when compared to lower VEGF concentrations and all VEGF concentrations present within alginate biomaterial. Error bars = SD. One-way ANOVA analysis performed with Tukey’s post-hoc test applied for multiple comparisons. *, **, *** and **** denotes that staining of blood vessel marker, anti CD31 shows a significant increase (p = <0.05, <0.01, <0.001 and <0.0001 respectively).

In parallel with increases in blood vessel infiltration in Laponite gels, haematoxylin and eosin (H&E) staining of day 21 Laponite and alginate samples containing 40 µg/mL VEGF showed differences in cell invasion. Laponite samples appeared rich in cellular content with regions of biomaterial still present whereas alginate samples contained large open spaces devoid of cells (Figure 4A). Auramine O, a substituted diphenyl-methane cationic dye, was used to label implanted nanoclay against biological tissue on the basis of a dramatic increase in the fluorescence emission upon stabilisation by clay nanoparticles [33,34]. A substantial volume of nanoclay remained at 21 days but was fragmented with cells and tissue present throughout the fragments (Figure 4A).

**Figure 4.**
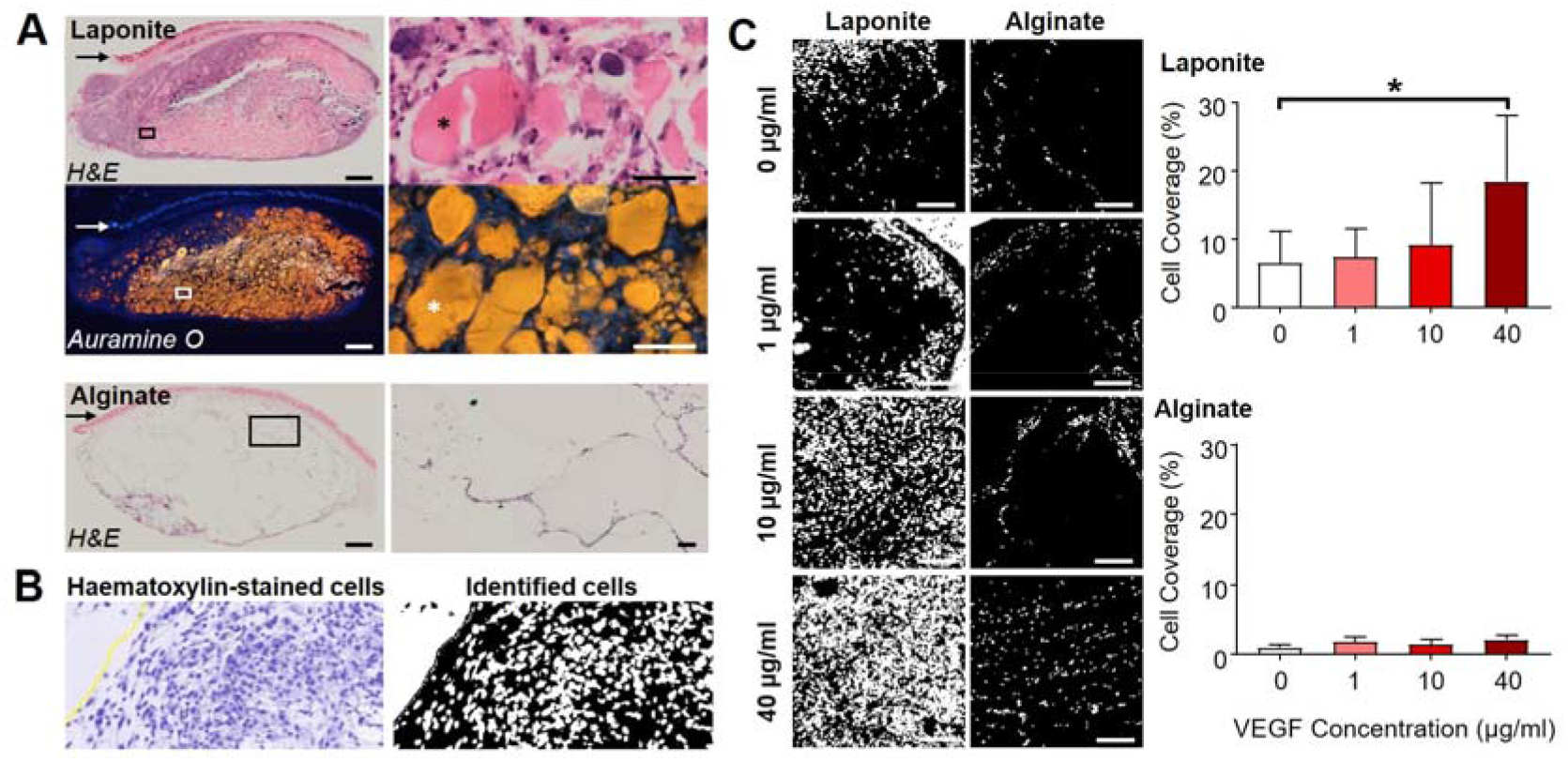
Cells invade and form blood vessels within Laponite in vivo. (A) Haematoxylin & eosin and Auramine O revealed pronounced cellular invasion in subcutaneously-implanted Laponite gels, with minimal cell invasion in alginate scaffolds. Left panel scale bar = 1 mm, right panel scale bar = 200 µm. Arrows indicate the skin epithelium; * indicates residual nanoclay gel. (B) Example image of high powered magnification of cells stained with haematoxylin (left panel); using computer-based software, cell invasion into biomaterial was identified, segmented and density of cells measured by percentage cell coverage. Scale = 100 µm. (C) Segmented cell analysis showed greater cell invasion in Laponite-treated groups when incorporated with increasing VEGF concentrations; in contrast alginate treatments showed very minor changes. (D) Quantification of cell coverage as described in (C) confirms this trend; 40 µg/ml VEGF mixed within Laponite exhibited a significant difference, no trend was observed for alginate. Error bars = SD. One-way ANOVA analysis performed with Tukey’s post-hoc test applied for multiple comparisons. * denotes that p = <0.05.

Analysis of H&E images involved software-based segmentation of haematoxylin hotspots to identify cell nuclei and determine gross cellular infiltration (Figure 4B). Image analysis showed that there was a trend of greater cellularity in Laponite samples with respect to increasing VEGF concentrations, with significantly greater cell infiltration at 40 µg/ml (Figure 4C, *p* = < 0.05). In contrast, there was no significant difference in cell infiltration for alginate gels irrespective of VEGF concentration (Figure 4C, 0 vs 40 µg/mL alginate, *p* = 0.14).

## Discussion

This study demonstrates the utility of a robust, well defined biomaterial in sequestering and preserving the activity of VEGF over prolonged periods of time *in vivo*. We showed that VEGF can be stably incorporated into injectable Laponite nanoclay gels, that the VEGF was active both *in vivo* and *in vivo*, and that the material acted as a stable template for angiogenesis and cell infiltration.

A medium concentration of 40 ng/ml of VEGF was necessary to promote maximum tubulogenesis in HUVECs cultured on Laponite substrates. This concentration is equivalent to the *total* VEGF required to elicit maximal tubulogenesis when pre-adsorbed to the Laponite surfaces (i.e. preincubation of Laponite substrates with 300 µL of 40 ng/ml medium (equal to 12 ng/cm^2^) before culture of cells on the surface resulted in the *same* degree of tubulogenesis as when 40 ng/ml was included in the medium concurrent with cell growth; Supplementary information 4 [S4] and [24]). This indicates that Laponite surface binding/interaction does not reduce the bioactivity of VEGF.

Relatively higher concentrations of VEGF were required to promote maximal tubulogenesis in HUVECs when delivered pre-mixed in Laponite. On first consideration, this might suggest that mixing reduces the activity of VEGF. While we cannot entirely exclude some loss of activity through mixing, it is important to recognise the qualitative differences in the approaches. In our study, 150 µL of Laponite gel was required to ensure a reproducible coating of each well (1 cm^2^ in area), equivalent to a mean depth/thickness of 1500 µm. By simple arithmetic it can be calculated that, at 5 µg/ml, the uppermost 24 µm of the ‘cylinder’ of Laponite would contain 12 ng VEGF per cm^2^ — the density of VEGF necessary for maximal tubulogenesis. Since, firstly, the approximate diameter of endothelial cells is 13-25 µm [35–37], secondly that there was a lack of any significant penetration of cells into the Laponite substrate *in vivo*, and thirdly that Laponite gels tightly sequester VEGF (Figure 1A), we consider it very likely that majority of VEGF in this ‘bulk mixed’ configuration is sequestered away from the cells and therefore is not bioavailable during the tubulogenesis assay.

Our *in vivo* findings, suggesting a localised, prolonged angiogenic effect of VEGF when sequestered in Laponite as compared with an alginate carrier that facilitates burst release, have intriguing corollaries in drug delivery. Conventional hydrogel drug delivery strategies tend to target controlled, sustained release from the biomaterial over time as necessary strategy for recruiting local endogenous cell populations [27]. Conversely, our data suggest that a sustained localised concentration of VEGF with rather minimal release is effective for promoting strong local angiogenesis. This observation is notable for two reasons. First the data implies that Laponite gels can host and even promote the invasion of responsive populations into the gel structure prior to a VEGF mediated response. A second implication is that, the activity of the sequestered VEGF is sustained in the gel before becoming bioavailable to invading cell populations.

VEGF has been shown to have a half-life of ≤ 90 minutes *in vivo* [38] and ∼34 minutes *in vivo* [39]. The observation of significant blood vessel formation from a single dose of VEGF 21 days after administration (a clear increase was apparent only between day 14 – 21), implies, either that the signalling important for subsequent vessel formation occurs at very early time points, or that the activity of VEGF is preserved and prolonged by incorporation in Laponite. Considering the first point, given a conservative estimation of a ∼90 minute half-life for VEGF (which is its known *in vivo* half-life in buffer at room temperature [38]) the quantity of active protein remaining from a 2 µg dose after only 24 hours would be at a sub-picogram range, whereas ng/ml concentrations are required for receptor activation *in vivo* [40,41]. Furthermore, cell invasion is likely to require time on the scale of days rather than hours and therefore at early time points (< 24 hours) initial presentation of VEGF would be limited to the biomaterial surface. Supporting this, at 21 days, penetration of Laponite by cells was incomplete, with cell-free areas visible in central regions of the nanoclay, suggesting gradual cell invasion. In addition, after implantation, cell invasion is likely to be initially mediated primarily by immune cells, such as neutrophils and macrophages, rather than the target cell population responsible for blood vessel formation (endothelial cells) as seen with other reports of biomaterial implantation [42,43]

Such considerations are underscored by the requirement in previous studies for much higher doses of soluble VEGF to be necessary to evoke a biological response, and data indicating that VEGF half-life can be extended by immobilisation in scaffolds. For example, a study by Galiano *et al.* on wound healing [44] showed it was necessary to add 20 µg of VEGF topically to each wound every other day for 10 days to exhibit a biological effect (a dose of 100 ug in total, compared with 2 µg total dose in the data presented in the current study). Furthermore it was observed that locally-applied VEGF induced undesired angiogenic responses at sites distant from the site of application, again suggesting that both protein degradation and protein diffusion are important factors for sustained, local activities. Other studies have sought to localise VEGF by passively binding it to scaffolds such as fibrin and have shown that, when coupled in this manner, VEGF can retain activity for periods of up to 2 weeks [45], again supporting the notion that Laponite may also stabilise VEGF (although perhaps by a different mechanism). There remains a paucity of data on how VEGF protein activity is stabilised by binding to other proteins or to biomaterials, but this remains an important area for future investigation. In view of these observations, we consider it likely that Laponite preserves the bioactivity of VEGF over a prolonged time *in vivo*.

Our approach has some limitations. For example, although we were able to measure the degree of blood vessel infiltration quantitatively by macroscopic and microscopic means in the presence of VEGF, we did not address the morphology of the vessels. It has been shown previously that sustained and high levels of VEGF_165_, can cause formation of dysfunctional blood vessels with irregular and leaky lumina [46–48]. Other suggest this may be due to subtle fluctuations in microenvironmental VEGF concentation [47]. We did not test directly the homogeneity of VEGF dispersion throughout the Laponite, but future studies may address this directly. Due to the highly sorptive nature of Laponite, we consider it likely that tight binding to nanoclay particles should provide precise control over microenvironmental concentration, however.

In contrast to other approaches in the literature (which often rely on complex bio-engineering of either the matrices from which growth factors are delivered [49] or the growth factors [50], or a combination of both [51]), our approach allows effective localisation of an active growth factor in an injectable hydrogel without complex chemical crosslinking methods. Since it is known that clay biomaterials also preserve the activity of another growth factor, bone morphogenetic protein 2 (BMP2) [25], exploration of nanoclay for delivery of a range of therapeutic proteins may be a rich area for future research.

## Acknowledgements

We acknowledge funding support from the EPSRC (EP/K018150/1 and Fellowship to JID), British Skin Foundation (ref S709), and NovoNordisk. DJP was supported by a Grundy Educational Trust Award. We acknowledge the dedicated and support and care of the staff in the Biomedical Research Facility, UoS, in particular Mike Broom, Andrew Crocker and Les Lawes.

