## Supplemental Information 1-4 for "Injectable nanoclay gels for angiogenesis"

### Supplementary Information for ‘Injectable Laponite clay gel for angiogenesis’

#### S1. Image Analysis of HUVEC 2D Tubule Network

The settings that were applied to compare tube networks were as follows: *Show maps of elements (single analysis)*, *show segments*, *Show nodes and junctions*, *Show meshes*, *Analyse master tree, Show extremities, Show branches, Show master segments* and *Show supressed isolated elements* was all checked. Default sizes were used but an *Iteration number* of 2 was applied. Phase contrast images were first converted to 8-bit by using the *Image/Type* module and then finally converted to red, green, blue (RGB) (Macro requires 8-bit RGB image for phase contrast analysis). The *Batch Image Treatment Tool* was used to analyse a batch of captured images using the phase contrast module. The macro generates a network overlay of each analysed imaged where it has identified the HUVEC network and an output data file; the *Tot. branching length* (pixels) was used to determine network organisation. A total of 6 wells per test group were analysed. 3-6 images were taken from each well and the mean calculated; these individual means were then used to calculate an over mean for that test group.

Please refer to Supplementary Figure S1.1 for further details of tubule image analysis with an example of an overlay of the ‘Identified Network.

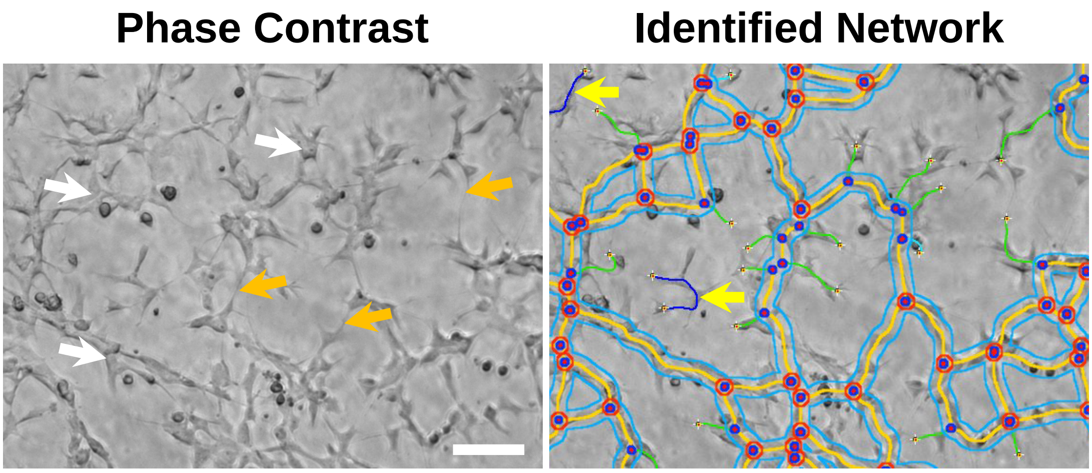


**Supplementary Figure S1.1.** Identified HUVEC Tubule Network by Image Analysis. (A) Image shows HUVECs cultured with 0.04 µg/ml VEGF in aqueous growth medium. Original phase contrast image shows individual HUVECs (black arrows) and tube extensions formed between adjacent cells (purple arrows). (B) Identified network using ImageJ ‘Angiogenesis Analyser’ plugin. Yellow lines depict ‘master segments of a continuous network (white arrows), with junctions (‘nodes) between cells identified by a red dot surrounded by blue circle (blue arrows). ‘Master junctions’ are ‘nodes’ surrounded by a red circle; these are major junction points that link 3 or more ‘master segments’ (red arrows). Green lines depict branch points of the network with terminal ends (extremities) and the light blue lines that surround ‘master segments’ are defined as mesh regions (‘loops’ of a network). Scale bars = 200 µm.

### S2. Subcutaneous Injection Study Questionnaire

#### Angiogenesis Scoring of Laponite Biomaterial +/- VEGF

Examine all the images below. After examining all the images, go back to the beginning and then score each image category below based on the degree of vascularisation from 0-6 (see below for more detail):

**Categories:**

BMS = Biomaterial score --> The degree of vascularisation (e.g. redness, blood vessels) **within and/or around the edge of the biomaterial.**

**0* = no blood vessel perfusion within biomaterial**
**2* = slightly red within biomaterial OR at the periphery**
**4* = biomaterial appears very red in colour with evidence of some blood vessel sprouting**
**6* = Significant blood vessel perfusion; biomaterial is totally red or dark red, lots of mature vessels growing within.**

GIS = Gross Image score --> The degree of vascularisation (e.g. redness, blood vessels) **within the whole image frame (biomaterial and surrounding tissue).**

**0* = No blood vessels in whole image frame (biomaterial or surrounding tissue)**
**2* = Some mature blood vessels/redness present in whole frame (biomaterial or surrounding tissue)**
**4* = Evidence of lots of blood vessels/redness within biomaterial and/or surrounding tissue**
**6* = Massive blood vessel perfusion within the biomaterial and lots of vessels around the outside.**
***Use 1, 3 or 5 scoring for images that do not quite match these scoring groups (in-between values)**

**Useful Information**

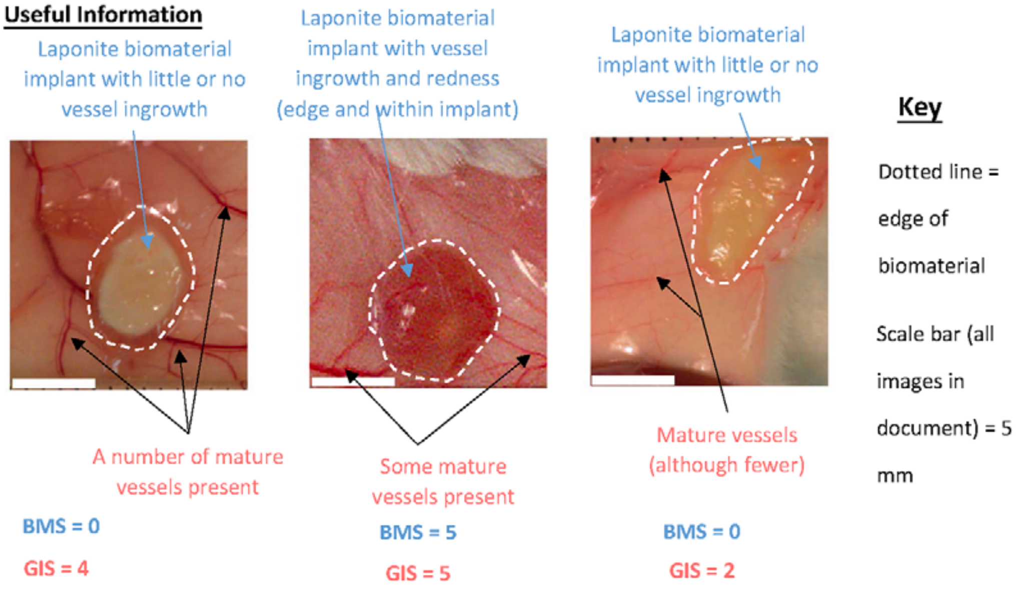

#### S3. Orbit Image Analysis Software Settings for Cellularity Dataset

#### General Configuration Settings

*Feature configuration Settings:*

Within the *Feature Configuration* window, the following settings within the *Classification* tab were applied:

*Structure size:* 4*; Median filter radius:* 0*; RGB channels used:* all checked*; Color Deconvolution Setup:* H&E*; Color Deconvolution Staining:* Stain 1 checked*; Classes for retrieving features/histograms:* leave blank*; Active fluorescence channels:* leave blank.

Within the *Segmentation* tab the following settings were applied:

*Set Features for Secondary Segmentation:* leave blank; *Cytoplasma Segmentation:* leave blank; *Minimum segmentation size:* 10; *Maximum segmentation length:* 500; *Minimum open distance:* 3; *Segmentation scale factor:* 1.0; *Disable object splitting:* unchecked; *Combine cross tile objects:* unchecked; *Discard tile border objects:* unchecked; *Mumford-Shah segmentation (cell clusters): Obj size* – 18 pixels, *Intens split* – 5; *Dilate:* 1; *Erode:* 0; *Dilate before erode:* checked; *Despeckle:* 0; *Smooth objects (GraphCut):* 0.0; *Nerve Detection Mode:* unchecked.

Within the *ROI* tab the following settings were applied:

*Use annotations as ROI:* All Groups; *Fixed circular ROI:* 0 pixels; *ROI offset X:* 0; *ROI offset Y:* 0.

Within the *Image Adjustments* tab the following setting was applied:

*Use image adjustments:* unchecked.

*Configure Classes Settings:*

Within the *Class Configuration* window (click button or press F4 on keyboard), two classes were defined: (1) *Background* in purple and (2) *Celltype 1* in white. Both classes had the *Only if used in Exclusion Model* option left as undefined.

**Classification Model Training**

The *Classification* module tab was selected and trained using a selection of ≥8 different images. As a rule, 1-2 images per sample group were chosen, there were 8 different treatments groups (4 x Laponite, 4 x alginate) so a minimum of 8 images were used to train the model.

After loading selected images, the *Background* class was selected (within F4 menu). Within the *Classification* module tab, the *Polygon* draw tool was selected and used to trace around select regions which best represented the background. This included areas that had no staining and regions where cells were not stained (e.g. wound granulation, healthy skin tissue matrix). After 6 regions had been traced, the second class (*Celltype 1*) was selected within the *Background* class menu. Again, the *Polygon* tool was selected and traced around regions where there were cells present. In total, 6 cells or region of cells were traced. Following this, the same set of steps were repeated with all other selected images.

After image tracing, the *Train* option (F7 on keyboard) within the *Machine Learning* module was selected to train the software to store pixel information regarding background and stained cells based on the region of interest (ROIs). Any models that were trained were then saved for future batch use.

**Region of Interest (ROI) Selection**

To analyse an image using a trained model, an ROI was selected using the *Annotations* module tab (right-hand side of the main program window). To do this, the *add polygon* tool was selected to trace around an ROI (e.g. the wound granulation tissue). After the ROI(s) had been drawn, they were selected in the *Annotations* module and the *edit annotation* option selected; within this window the *Type* field was changed to *ROI*. Once and ROI had been defined this information is automatically stored within the *OrbitOmero.properties.template* file located within the user profile of the program (please refer to manufactures instructions for more details). Therefore, images can be closed and re-opened for future analysis.

**Image Analysis (Single)**

After loading an image and training a model, images can be analysed for classified areas. To use a pre-existing model, it was loaded via the *Model* tab module. Then within the *Classification* module tab, the *Classify option* was selected (F8 on keyboard). The resulting output contained the ratio of both the *Background* class and *Celltype 1* class. Values generated from the *Celltype 1* class were multiplied by 100 to determine the % coverage of pixels that were identified as cells.

**Image Analysis (Batch)**

Most of the cellularity data was generated by analysing a batch of images following model training. First, a pre-existing trained model was loaded and then the *Batch* module tab selected. Using the *Local Execution* option, multiple files could be selected for analysis using the class attributes within pre-loaded model. The output window contained the ratio of both the *Background* class and *Celltype 1* class. Values generated from the *Celltype 1* class were multiplied by 100 to determine the % coverage of pixels that were identified as cells.

**Supplementary Information 4**

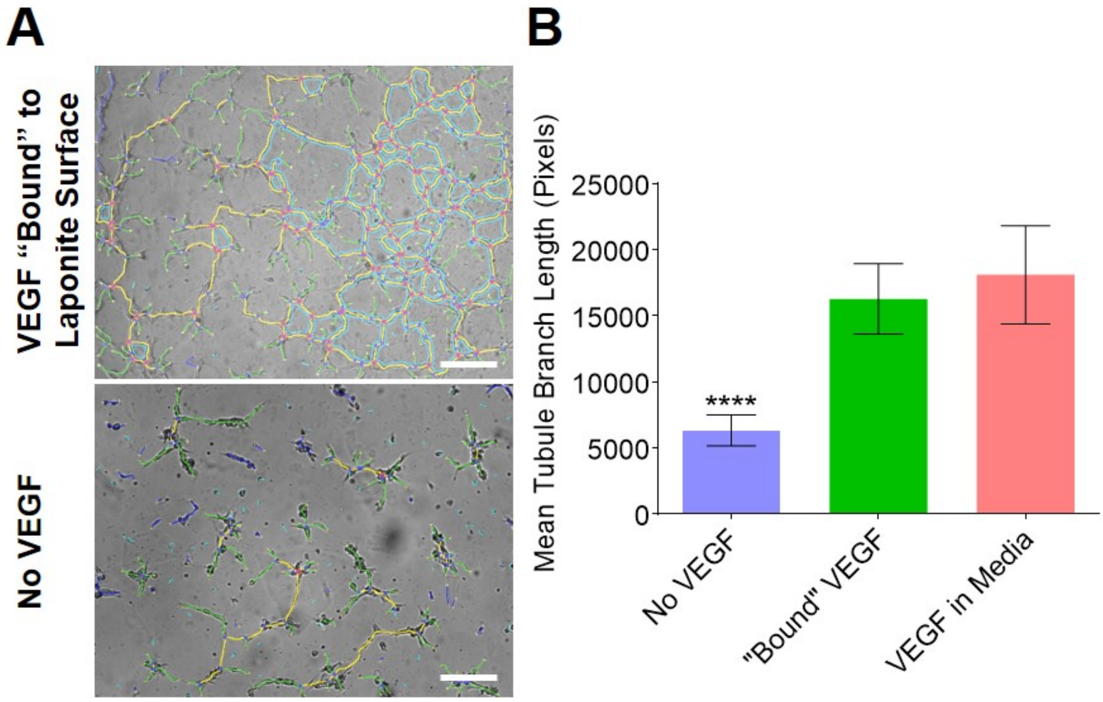


VEGF surface adsorbed to Laponite gels elicits an equivalent increase in cell tubule formation to VEGF incorporated in medium. Refer also to Figure 1.
